# The precedence effect in spatial hearing emerges only late in the auditory pathway

**DOI:** 10.1101/2021.07.21.453295

**Authors:** Kongyan Li, Ryszard Auksztulewicz, Chloe H. K. Chan, Ambika Prasad Mishra, Jan W. H. Schnupp

## Abstract

**Background:** To localize sound sources accurately in a reverberant environment, human binaural hearing strongly favors analyzing the initial wave front of sounds. Behavioral studies of this “precedence effect” have so far largely been confined to human subjects, limiting the scope of complementary physiological approaches. Similarly, physiological studies have mostly looked at neural responses in the inferior colliculus, or used modeling of cochlear mechanics in an attempt to identify likely underlying mechanisms. Studies capable of providing a direct comparison of neural coding and behavioral measures of sound localization under the precedence effect are lacking.

**Results:** We adapted a “temporal weighting function” paradigm for use in laboratory rats. The animals learned to lateralize click trains in which each click in the train had a different interaural time difference. Computing the “perceptual weight” of each click in the train revealed a strong onset bias, very similar to that reported for humans. Follow-on electrocorticographic recording experiments revealed that onset weighting of ITDs is a robust feature of the cortical population response, but interestingly it often fails to manifest at individual cortical recording sites.

**Conclusion:** While previous studies suggested that the precedence effect may be caused by cochlear mechanics or inhibitory circuitry in the brainstem and midbrain, our results indicate that the precedence effect is not fully developed at the level of individual recording sites in auditory cortex, but robust and consistent precedence effects are observable at the level of cortical population responses. This indicates that the precedence effect is significantly “higher order” than has hitherto been assumed.

## 1 Introduction

Having two healthy ears can bring considerable advantages, enabling us to use binaural cues to localize sound sources and to separate sound sources in a cluttered acoustic environment. However, sound localization is often rendered more complicated by the fact that every hard surface reflects sound waves, acting like a “sound mirror” and creating a mirrored sound source which interferes with the localization of the original source. Interestingly, we are usually largely unaware of the powerfully distorting effect such reverberation has on the sound waves that arrive at our eardrums. Our auditory pathways appear to be equipped with powerful “echo suppression” mechanisms, but their function and their physiological basis remain very poorly understood. One important part of this echo suppression which has been studied in some detail is the so-called “precedence effect”. This refers to the phenomenon that the perception of sound source direction puts great emphasis on the binaural cue values experienced during the first few milliseconds of a new sound burst. This reliance on sound onset cues is thought to reduce the confounds that can arise when the binaural cue values of later parts of the sound are distorted by interference from reverberant sound (for a review, see Litovsky et al., (1999)).

The precedence effect has been recognized as a psychophysical phenomenon in humans for over 100 years (for a history see Gardner, (1968)), and behavioral studies on rats (Hoeffding, Harrison, 1979) and cats (Cranford, Oberholtzer, 1976) indicate that it may be a common feature of mammalian hearing. However, previous animal work tended to use brief individual stimulus pulse-echo pairs delivered in the free field, which is fine for investigating whether or not there is a precedence effect, but it does not permit quantification of the relative weighting that the precedence effect might give to various elements of a stimulus composed of a rapid series of consecutive pulses. Understanding the perceptual weighting in pulse train stimuli has considerable potential ecological and translational importance, firstly because the majority of animal vocalization sounds, including human vowels, are effectively (band-pass filtered) pulse trains, and secondly because cochlear implant (CI) neuroprosthetic devices which are used increasingly for the treatment of profound to severe hearing loss, will typically encode auditory stimuli as rapid trains of electrical pulses delivered to auditory nerve fibers. Being able to measure the relative perceptual weight that the brain assigns to each pulse in a pulse train as it forms sound source location judgments is therefore likely to become important in the development of better sound processing algorithms for future assistive or neuroprosthetic devices.

One elegant way to perform such a quantification was introduced by Stecker and Hafter, (2002), who used brief binaural click trains delivered over headphones to measure so-called temporal weighting functions (TWFs) psychoacoustically. Each click in the train carries its own binaural cue parameters (interaural time or level differences, ITDs or ILDs), but clicks are delivered at a rate high enough for clicks to fuse perceptually into a single perceived sound. Subjects are asked to indicate the perceived source direction of the click train, and a statistical analysis is used to calculate the relative influence of each click in the train on the overall perceived source direction. Studies using this paradigm on normally hearing human subjects (Brown, Stecker, 2010; Brown, Stecker, 2011; Stecker et al., 2013; Stecker, 2014) have consistently shown a strong onset dominance to click trains, which is more pronounced for higher click rates than for lower click rates.

From a translational perspective it would be of interest to extend this type of approach to hearing impaired patients as well as to animal models suitable for the study of hearing loss and treatments. It is well known that patients with severe hearing loss often have great difficulty processing binaural cues, even if they are fitted with bilateral CIs (van Hoesel, Tyler, 2003). The processing of ITDs appears to be particularly severely affected in such patients, even more so than that of ILDs, and in this study we shall focus on ITDs as they are the type of binaural cue for which deficits, and therefore the need for improvement, are greatest. Some recent studies have raised the intriguing possibility that changes to the current standard algorithms for computing CI stimulus pulses might enhance the ITD sensitivity of bilateral CI patients. For example, introducing temporal jitter into the interval between stimulus pulses may enhance the sensitivity to ITDs in bilateral CI users (Laback, Majdak, 2008) as well as normally hearing listeners (Goupell et al., 2009). Meanwhile, Litovsky, (2006) reported that bilateral CI users and normally hearing children may not exhibit a precedence effect, which raises questions about how stimulation paradigms and neurobiological development interact to further result in effective spatial auditory perception. We will need a better understanding of the technical factors and biological mechanisms that determine binaural listening performance if we want to improve patient outcomes, and the development of a suitable animal model which lends itself to combined behavioral and physiological approaches is going to be an important step in this endeavor.

Our recent papers (Li et al., 2019) demonstrated that Wistar rats have similar sensitivity to both onset and ongoing envelope ITDs as humans when tested in a sound lateralization task with pulsatile stimuli, and, intriguingly, rats fitted with bilateral CIs can exhibit essentially normal behavioral ITD sensitivity even after prolonged neonatal hearing loss (Rosskothen-Kuhl et al., 2021). Rats may thus be a highly valuable model for the study of binaural hearing if they also exhibit similar temporal weighting as human subjects do. In this study we therefore sought to answer the following three questions: 1) whether it is possible to adapt the TWF approach developed by Brown and Stecker, (2010) for behavior testing in rats, 2) whether behaviorally measured ITD TWFs for rats resemble those reported for humans, with most or all of the perceptual weight vested in the first click of a train, and 3) whether or how electrophysiological responses recorded from rat auditory cortex reflect the behavioral weighting.

## 2 Materials and Methods

### 2.1 Animals

Our subjects were nine female Wistar rats which were 8-week-old and weighed 216-242 g at the beginning of the study. Of these, four were first used for behavioral training and psychoacoustic determination of ITD TWFs (see 2.2). All nine (4 trained and 5 naive animals) were then used in terminal electrophysiological experiments to elucidate the cortical encoding of the binaural stimuli (see 2.3). The rats were housed in standard cages with 2 or 3 rats in each.

Preyer’s reflexes were tested and the outer ears and tympanic membranes were visually examined to ensure that the rats had healthy, sensitive hearing. In addition, prior to the behavioral and electrophysiological experiments, acoustic brainstem responses (ABRs) were recorded. For this examination, the rats were anesthetized by intraperitoneal injection of ketamine (80mg/kg, 10%, Alfasan International B.V., Holland) and xylazine (12 mg/kg, 2%, Alfasan International B.V., Holland). Eye gel (Lubrithal, Dechra Veterinary Product A/S Mekuvej 9 DK-7171 Uldum) was applied to prevent the eyes from drying. The outer ear canals and tympanic membranes were inspected under microscope (RWD Life Sciences, China). The rats were then fitted to a stereotactic instrument with a pair of hollow ear bars in a sound attenuating chamber, and ABRs to clicks were recorded to ascertain low hearing thresholds in both ears.

### 2.2 Behavioral study

#### 2.2.1 Behavioral training setup

The behavioral setup and the training methods were identical to those described in sections 2.2 and 2.3 of Li et al., (2019). In brief, a training box was situated in a sound attenuating box and the front wall of the training box was fitted with three brass water spouts. Two hollow tubes were connected to a pair of mini headphone drivers (GQ-30783-000, Knowles, Itasca, Illinois, US) to deliver the sound stimuli delivered by a USB sound card (StarTech.com, Ontario Canada, part No. ICUSBAUDIOMH) and amplified by an audio amplifier (Adafruit stereo 3.7W class D audio amplifier, part No. 987) into the behavioral training box as close to the rats’ ears as possible. Stimulus delivery and monitoring and control of the behavioral task were performed by a Raspberry Pi computer running custom written Python software.

#### 2.2.2 Behavioral training task

During behavioral training and testing, the animals were tested five days a week, with two rest days. Our training uses drinking water as a positive reinforcer. Therefore, a day prior to the first testing day, the home cage water bottles were removed, and for the following 5 days, the rats only had access to drinking water during their twice daily testing sessions. They then had easy access to *ad lib* water from the evening of the fifth training day until the morning of the second rest day. Food was available *ad lib* in the animals’ home cages throughout.

Behavioral training and testing were essentially identical to Li et al., (2019), except that a different set of stimuli was designed and used to enable the quantification of TWFs. In the behavioral experiments, rats performed a two-alternative forced-choice (2-AFC) near-field lateralization task. Rats initialized each trial by licking a centrally positioned “start spout”. Initiating a trial was rewarded with a small drop of water on a random subset of 1 in every 7 trials. Initiating a trial triggered the delivery of a binaural stimulus, to which the animals responded by licking one of two “response spouts” positioned either side of the start spout. If the animal’s choice corresponded to the side indicated by the binaural cues, it was rewarded by three small drops of water delivered through the response spout. If the response was incorrect, it triggered a 15 s “timeout” during which a 90 dB negative feedback sound was played and no new trials could be initiated. If the rat made a wrong response, the following trial would be a “correction trial”, in which the last stimulus was repeated. “Correction trials” help reduce the tendency of animals to develop responses biases towards one side, but are excluded from the calculation of the correct response scores. Each rat performed two sessions per testing day, one in the morning and one in the afternoon, each session lasting ∼20 minutes. The animals would typically perform between 100 and 200 trials per session.

The rats were initially trained with 200 ms long, 300 Hz binaural click train stimuli which contained both ILD (+/- 6 dB) and ITD cues (+/- 0.136 ms). Our convention here is to use negative values to indicate binaural cues that favor the left ear. We required that the rats lateralized these initial training stimuli at least 80% correct in at least 2 sessions to advance to the next “ITD-only” training stage, during which ILDs were set to 0 dB. The rats reached the 80% correct criterion in this first stage of training after 8-10 days of training. After the initial training phase, all stimuli presented throughout the rest of the study had 0 dB ILDs and varied in ITD only. Once the rats reached 80% correct on two sessions with the 0.136 ms ITD only stimuli, we increased the range of ITD values tested in each session. During this stage, the ITDs presented at each trial were drawn at random from the set ±[0.1587, 0.136, 0.0907, 0.068 ms, 0.0454 ms, 0.0227] ms. This set purposefully includes some ITD values that are below previously determined perceptual ITD thresholds for rats (∼0.05 ms, see Li et al., (2019)), in order to accustom the animals to the possibility that sessions may include trials that may be difficult to lateralize. In this potentially more difficult “wide ITD range” training stage, the rats had to reach 75% correct in at least two sessions before advancing to the TWF testing stage. Animals which did not quickly advance to that final stage were given additional training sessions with easier stimuli during which timeouts and reward quantities were individually adjusted as necessary to achieve reliably high levels of performance.

#### 2.2.3 Acoustic stimuli for behavioral TWF measurement

The stimuli used in the TWF measurement phases of our experiments were modeled on the stimuli developed by Brown and Stecker, (2010). Our stimuli consisted of trains of 8 binaural clicks (**Figure 1**) delivered at rates of either 20, 50, 300 or 900 Hz. Importantly, the ITD of each click in the train was varied by introducing a small, random “temporal jitter” in the timing of each click which was independent in each ear. An analysis of lateralization judgments for a large number of stimuli with different ITD values at each click in the series would then make it possible to determine the “weight” (that is, the contribution made) by the n-th click in the train to the perceived lateral position of the click train as a whole.

**Figure 1:**
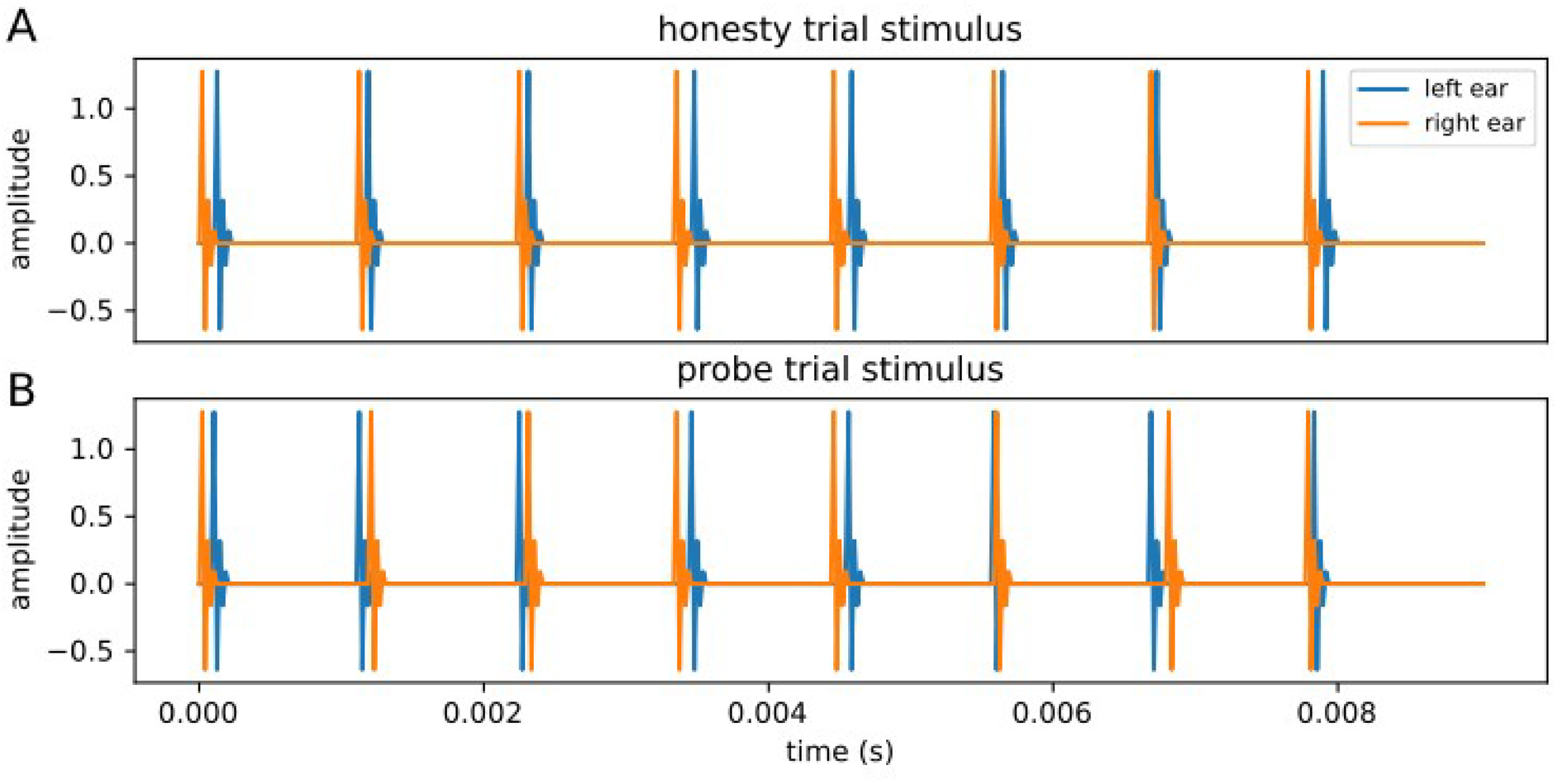
Example waveforms of the acoustic stimuli used in the behavioral experiments. **A**: Example of an honesty trial stimulus at 900 Hz with 0.042 ms jitter and +0.083 ms offset (+: right ear leading, –: left ear leading). The time differences between the two clicks in each click pair were in the range of +0.042 ms to +0.125 ms, which means every click pair leading to the right ear. There is no ambiguity and the rat will only be rewarded for responding on the right. **B**: Example of a probe trial stimulus at 900 Hz with 0.125 ms jitter and 0 ms offset. The time differences between the two clicks in each click pair were randomly chosen from the range of – 0.125 ms to + 0.125 ms. Since there was no *a priori* objectively correct response to probe trials, rats were rewarded in those trials irrespective of which response spout they licked.

A potential difficulty with the use of these stimuli is that one cannot always determine *a priori* whether a subject’s lateralization response is “objectively correct”, because ITD cue values of different clicks in the train could, by design, point in opposite directions, and how such ambiguous stimuli with conflicting ITD values are “supposed to be perceived” depends on the subject’s own TWF which is unknown at the start of the experiment. Brown and Stecker, (2011) simply trusted their human participants to understand the objective of the experiment and to report their lateralization judgments faithfully, presumably with low error or bias, and without the need of trial-by-trial reinforcement. Our rats, in contrast, need to be kept motivated and honest throughout the experiment by regular rewards for “correct” responses. Therefore, we constituted each block of trials as a randomly interleaved set of “honesty trials” and “probe trials”. In honesty trials, the ITDs of all clicks in the 8 click sequence pointed in the same direction, that is, they were either all positive (right ear leading) or all negative (left ear leading), so that the response to these honesty trials could be judged objectively as correct or incorrect irrespective of the details of each animal’s TWF. Clicks in honesty trails had a fixed ITD offset of ± 0.083 ms, plus an additional jitter drawn uniformly at random from a range of ± 0.042 ms, in steps of 10.4 μs afforded by the 96 kHz sample rate Hi-Fi USB Audio sound card. Since most ITD values in an honesty trial should be above typical rat ITD thresholds reported in Li et al., (2019) we expected them to be relatively easy to lateralize correctly, and responses to honesty trials were only rewarded if the animal responded on the appropriate side. We required that the rats lateralized at least 80% of honesty trials correctly in at least two sessions before they would also receive “probe trials”, in which the ITD for each click was drawn independently and uniformly from the range of ± 0.125 ms and ITDs of subsequent clicks in the train were allowed to point in opposite directions. Responses to probe trials were always rewarded regardless of the side on which the animal responded. In each TWF testing session, honesty trials, and probe trials were randomly interleaved at a ratio of 2:1. The large proportion of honesty trials ensured that random guessing without attending to the sounds was not an effective strategy for the animals and allowed us to monitor that the animals continued to report their lateralization percepts with good accuracy throughout. During informal testing, the authors were unable to distinguish honesty trials from probe trials just by listening to them, and there is no indication that the rats could distinguish these either. We therefore consider it safe to assume that the rats’ responses to the probe trials accurately reflected their lateralization judgments for these stimuli. After reaching the “ITD only” lateralization training criteria described above, all four rats were also able to meet the 80% correct TWF honesty trial criterion after minimal training, as might be expected given that to casual human observers, TWF stimuli with jittered ITDs and stimuli with fixed ITD sound indistinguishable.

#### 2.2.4 Behavioral data analysis

TWFs were computed from the responses to the probe trials only, and separately for each of the four click rates (20, 50, 300 and 900 Hz), by computing a Probit regression to fit the probability of a “right spout” response against the ITD values for each of the 8 click pairs in the train, using the open source Python function *statsmodels.discrete.discrete_model.Probit* (Seabold, Perktold, 2010). The Probit regression model takes the form:

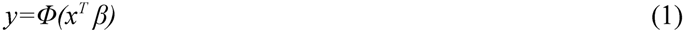

Here, *y* is the probability that the animal will respond on the right, *x* is the vector of the eight ITD values of the clicks in the train plus an added 1 for the intercept, and *β* are the coefficients, or “weights” attributed to each of the clicks, which are estimated by maximum likelihood. *Φ* is the cumulative Gaussian normal distribution. The fitted model thus assumes additive effects of the weighted ITD of each click in the train, and the set of coefficients *β* represent the animal’s TWF. Below we will use the terms Probit coefficient and temporal weight interchangeably.

### 2.3 Electrophysiology

#### 2.3.1 ECoG recording apparatus

In the electrophysiological experiment, the same four rats which performed the psychoacoustic testing were used. Acoustic stimuli were generated by RZ6 multi-I/O processor (Tucker-Davis Technologies, USA) and presented via a pair of custom-made speakers (AS02204MR-N50-R, PUI Audio, Inc.) fitted to the openings of the hollow stainless steel ear bars, which fixed the rat into a stereotactic instrument (RWD Life Sciences, China). The speakers were calibrated with a GRAS 46DP-1 microphone (GRAS Sound & Vibration A/S), and their transfer functions were compensated with an inverse filter to be flat over the range of 600 Hz to 20 kHz to ∼ +/- 3dB.

Neural activity was recorded using a 61-channel electrocorticographic (ECoG) array (Woods et al., 2015). The flexible (∼30 µm thin) ECoG array comprised 203 μm diameter circular electrodes arranged on an 8×8 square grid, with three of the four corner positions unoccupied, and a 406 μm spacing between neighboring electrodes. The array covered an area of 10.6 mm^2^.

The neural signal was captured through two Intan C3314 32 channel headstage amplifiers (Intan Technologies, USA) connected to a PZ5 neurodigitizer (Tucker-Davis Technologies, USA), and processed with RZ2 bioamp processor (Tucker-Davis Technologies, USA). Python programs written by the authors were used to generate stimuli and save the recorded signals.

#### 2.3.2 ECoG recording procedure

ECoG was recorded from the auditory cortex (AC). At first, rats were anesthetized as described in the ABR recording procedure in 2.1, and their scalp was shaved. Prior to ECoG recording, ABRs were tested again to make sure the ear bars were still in good position, followed by intraperitoneal injection of urethane (20%, 1mL). If a toe pinch reaction was observed during the ECoG recording, an additional 1 mL of urethane was injected. The total amount of injected urethane was less than 7.5 mL/kg. Additionally, butorphanol (10 mg/mL, 0.2mL/kg every 1-2 h, Richter Pharma AG, 4600 Wels, Austria) was subcutaneously injected as a painkiller during surgery. A deep cut in the midline of the scalp was made and the surgical field was exposed. Local anesthetic Lignocaine (0.3 mL, 20 mg/mL, Troy Laboratories Pty Ltd, Australia) was applied on top of the surgical area. A craniotomy was performed over the right, or, in most cases, both temporal cortices. From a point 2.5 mm posterior to Bregma, a line was drawn perpendicular to the sagittal suture to the temporal ridge, and the intersection of this line and the ridge was marked. The craniotomy area extended 5.0 mm posterior and 4.0 mm ventral from this intersection point, to allow the placement of an ECoG electrode array on the auditory cortex. A hole was drilled through the skull anterior to Bregma to fix a screw which served as a reference electrode to connect to the ground wire of the recording headstage amplifier.

After placing the ECoG electrode array on the AC, acoustic stimuli were presented to the rat. ECoG neural signals were recorded at 6 kHz sample rate. At the end of the recording experiments, the rats were euthanized with an overdose of Pentobarbital (1∼2 mL, 20%, Alfasan International B.V., Holland).

#### 2.3.3 “Sparse” TWF stimuli for electrophysiology

The ECoG recording experiments used acoustic stimuli which were similar to those used for behavior described in section 2.2.3, but simplified and optimized for estimating TWFs based on neural activity. Identifying the neurophysiological correlates of TWFs is more challenging than measuring TWFs behaviorally, since neurons in the auditory pathway do not give binary “left” or “right” responses, but instead have ITD tuning curves which can differ greatly from one neuron to the next. Furthermore, these tuning curves may map ITD values to neural firing rates in a non-monotonic manner (Fitzpatrick et al., 2000). Measuring neural TWFs can therefore easily degenerate into an under-constrained problem, where one seeks to understand the mapping of a relatively large number of continuous-valued stimulus parameters (the ITDs of each click in the train) onto a very noisy continuous-valued output variable (the neural firing rate) through a set of tuning curves of unknown shape. In an attempt to make the problem more tractable, we reduced the complexity of the stimuli compared to those used in the behavioral experiment by reducing the number of clicks in the train from 8 to only 4, and by constraining each click so that it could take only one of two possible ITD values, either -0.164 ms or +0.164 ms. These ITD values are close to, and slightly larger than, the top of reported +/- 0.13 ms normal ecological ITD range for rats (Koka et al., 2008). By constraining each stimulus click train to have only four clicks, and each click constrained to take one of only two possible values (corresponding to “far left” or “far right”), we reduced the set of all possible stimulus click trains to only 16, and we presented each of these 16 possible stimuli 40 times, in pseudo-random order, recording 640 responses in total at each recording site. We refer to this reduced-complexity set of stimuli as “sparse” TWF stimuli to distinguish it from the richer set of stimuli used in the behavior. The tested click rates were 300 Hz (for a total click train duration of 13.496 ms) and 900 Hz (for a click train duration of 4.608 ms).

#### 2.3.4 ECoG data analysis

We attempted two approaches to analyze the electrophysiological responses, one “univariate” approach which used a regression model to try to account for neural response amplitudes observed at each individual recording site, and one “multivariate” approach which attempted to use recently developed population decoding analyses to reconstruct stimulus ITDs from single trial population responses observed across the ECoG array. The multivariate approach turned out to be much more successful, which has interesting implications for the nature of the representation of perceived ITDs as we shall see below.

##### 2.3.4.1 Univariate analysis: channel-wise regression

Our analysis of the responses recorded with these stimuli was based on the assumption that most ITD sensitive neurons in the central auditory pathway would be tuned so as to have a “preference” for ITDs pointing to the contralateral side, while a minority might have an ipsilateral preference, but very few should have tuning curves that are symmetric at either end of the ecological range of ITDs (Benson, Teas, 1976; Woldorff et al., 1999; Yao et al., 2013). We further assumed that neural response amplitudes of contralaterally tuned units should consistently increase when contralateral leading ITDs are presented, irrespective of whether these contralateral ITDs occur at the first, second or n-th click. Similarly, for ipsilaterally tuned units, response amplitudes should consistently decrease when contralateral ITDs are presented. Under these simplifying assumptions we can attempt to fit TWFs to the neural data using a simple multiple linear regression, which regresses response amplitude against the signs of the four ITDs in each stimulus.

This univariate analysis of ECoG voltage data was further based on standard methods for quantifying evoked response amplitudes from LFPs as follows. First, per channel, the signal was band-passed using a 4^th^ order band-pass filter from 30 Hz to 300 Hz (scipy.signal.butter(), scipy.signal.filtfilt()). The band-passed signal was downsampled by a factor of 4 to a sample rate of 1500 Hz (scipy.signal.decimate()) and the decimated multichannel data were denoised using the “denoising by spatial filtering” methods developed by de Cheveigné and Simon, (2008). The cleaned data were “re-referenced” by subtracting the median across all channels (Liu et al., 2015). Neural responses were then quantified by epoching the cleaned, re-referenced signals into data segments ranging from 1 ms to 30 ms post stimulus onset. This epoch was chosen by visual inspection to cover the onset response peak, and it is consistent with reports that the initial responses to acoustic stimuli in the rat auditory cortex peak approximately 20 ms after stimulus onset (Rutkowski et al., 2003). The epoched data were baseline-corrected by subtracting their mean, and the RMS amplitude was calculated for each response epoch. Outlier epochs with RMS amplitudes greater than three standard deviations above the median RMS amplitude were excluded from further analysis.

The distribution of RMS response amplitudes obtained in this manner was highly positively skewed. To make it more suitable for linear regression analysis to obtain TWF values, we therefore log-transformed the RMS values. Furthermore, we wanted to compute temporal weighting coefficients in normalized units which were insensitive to site-to-site or animal-to animal variability in the range of observed voltage values that can result from variable electrode impedances or electrode placements. We therefore z-scored the log(RMS) values prior to regression analysis.

The transformed data were then subjected to an ordinary least squares (OLS) regression (statsmodels.api.OLS, (Seabold, Perktold, 2010)) with constant added (**Equation** 2). The form of the regression model is

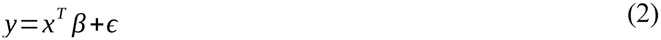

where *y* is the z-scored, log transformed LFP amplitude observed in each trial, *x* is a vector of the regressors (the ITDs of the 4 clicks in ms, and the added constant to provide the intercept), *β* is the vector of regression coefficients (the neural TWF weights in units of standard deviations of log RMS LFP amplitudes per ms of ITD), and *ε* is an error term which, as usual for normal linear regression, is assumed to follow a Gaussian distribution. In addition to computing the regression weights *β*, the software returned *p* values indicating how likely it is that the corresponding *β* is significantly different from zero.

##### 2.3.4.2 Multivariate analysis: population-based decoding

In addition to the mass-univariate (i.e., channel-by-channel) analyses described above, data were also subject to a multivariate analysis based on the response of the population of recorded neurons (i.e., pooling information from multiple channels). Rather than computing the “weight” of a given ITD in the train as scaling factor that maps ITD values onto changes in response amplitude, the rationale of this analysis was to try to decode the ITD value of each click in the train on a trial-by-trial basis from the pattern of neural activity measured by multiple ECoG channels and to quantify the “weight” of each ITD in the stimulus train by how well the ITD value can be decoded from single trial neural population responses using the methods originally described in humans by Wolff et al., (2017) and Auksztulewicz et al., (2019), and recently adapted for the rat auditory cortex (Cappotto et al., 2021). To this end, we first selected channels that showed a robust evoked response to the click train. The criterion that we used for channel selection was based on the signal-to-noise ratio (SNR), defined for each channel as the ratio between the RMS of the signal in the first 30 ms after click train onset and the RMS of the signal in the last 30 ms prior to click train onset. Only channels with SNR > 3 dB were taken into the analysis.

Following channel selection, for each ECoG array position, data from multiple channels were used to decode the ITD of each click in the train, one at a time, based on the RMS in the 0-30 ms time window following click train onset. To this end, we split the data into three sets: (1) the test trial itself, (2) the remaining trials with the same ITD value as the test trial, and (3) the remaining trials with a different ITD than the test trial. Based on these three sets, we obtained three vectors of average response RMS amplitude values concatenated across channels. We then calculated the multivariate Mahalanobis distance values between (1) the test trial vector and the average vector of trials with the same ITD, as well as (2) the test trial vector and the average vector of trials with a different ITD. The Mahalanobis distance values were scaled by the noise covariance matrix of all channels, i.e., the covariance based on single-trial residual RMS after removing the mean RMS from each trial (Muhle-Karbe et al., 2020). The scaled Mahalanobis distance values, obtained for a given trial *k* relative to other “same” or “different” trials, were used to calculate the overall decoding distance metric according to the following equation:

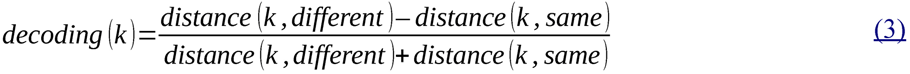

This procedure was carried out for each trial in turn in a leave-one-out cross-validation approach, and the resulting decoding values were averaged across trials to obtain ITD decoding estimates for each of the four clicks in the train. The decoding estimates were tested for statistical significance using a signed rank test. These signed rank tests were done for all electrode placements pooled together, as well as separately for 300 Hz and 900 Hz click rates. The tests were corrected for multiple comparisons using Bonferroni correction. Furthermore, to plot representative examples of individual electrode placements, we calculated 99% confidence intervals of the observed individual decoding estimates by repeating the analysis over 1000 iterations, whereby in each iteration the ITD labels were randomly reshuffled, to obtain a surrogate distribution of decoding estimates.

We also explored whether neural activity later than the first 30 ms following click train onset can be used to decode click ITDs. To this end, we repeated the decoding analysis in a sliding time-window approach, using a window length of 30 ms (with a time step of 5 ms). Specifically, for each time window, we extracted the RMS envelope (down-sampled to 200 Hz to yield 7 RMS values per time window), de-meaned it by removing the average across the time window (separately for each channel), and concatenated the de-meaned values across channels (Wolff et al., 2020). The resulting vectors of RMS fluctuations in multiple channels were used to calculate the Mahalanobis distance metrics and the corresponding decoding estimates, as described above. Decoding time series were tested for statistical significance for each click pair and time point using a signed rank test, correcting for multiple comparisons using a false discovery rate of 0.01 (Benjamini, Hochberg, 1995). Again, these statistical tests were applied for all electrode placements pooled together, as well as for 300 Hz and 900 Hz click rates separately.

## 3 Results

### 3.1 Behavioral task showed profound onset dominance

Using the protocols described above, the rats were trained twice daily, 5 days a week. Usually, the rats would perform ∼160 trials on average in one 20 minute training session, but the number could vary from just below 100, to well over 200. The rats were initially trained with combined ITD and ILD cues for 13 to 17 training sessions until they reached the 80% correct criterion in at least two sessions. They then progressed to “ITD only training”. The removal of the ILD cue had little effect on performance, so the rats required only 2 to 3 sessions to reach the criterion of being 80% correct in two sessions. They then progressed to multiple ITD value training, which introduced many new stimuli with smaller ITDs closer to threshold, and they needed between 18 and 21 training sessions before reaching the respective criterion for this training phase (75% correct in two sessions). We initially trained 5 rats with this protocol, four of which reached the required high performance with ITD-only stimuli after about two weeks of training. The one rat which failed to achieve the performance criterion for progression after 2 weeks of training was excluded from the rest of the study.

After completing the multiple ITD values training, the rats were trained with “honesty trial” TWF stimuli sessions. Once their performance was over 80% correct in two or more sessions, they were tested in TWF stimulus sessions containing both “honesty” and “probe” trial stimuli in a 2:1 ratio. For the 50 Hz and 300 Hz click rates, only 2 sessions were needed for all the rats to reach the final “honesty + probe” testing stage. For the 20 Hz and 900 Hz click rates, 4 and 6 sessions respectively were required to reach the final behavioral testing stage.

In the final, psychoacoustic testing stage, for stimulus pulse rates of 20 Hz, 50 Hz, 300 Hz, and 900 Hz, we collected data over a total of 35 sessions for rat 1801, 34 sessions for rat 1802, 33 sessions for rat 1803 and 33 sessions for rat 1805. The numbers of probe trials and honesty trials, and the correct rate in honesty trials in each condition for each rat is summarized in **Table 1**. Rat 1802 was the best performer in honesty trials for all click rates. The correct rate in honesty trials and the number of training sessions needed in the “honesty” training stage suggest that the task difficulty was similar across the four tested click rates. Note that even at the most difficult, 900 Hz click rate, all rats had more than 80% correct responses in honesty trials.

**Table 1:**
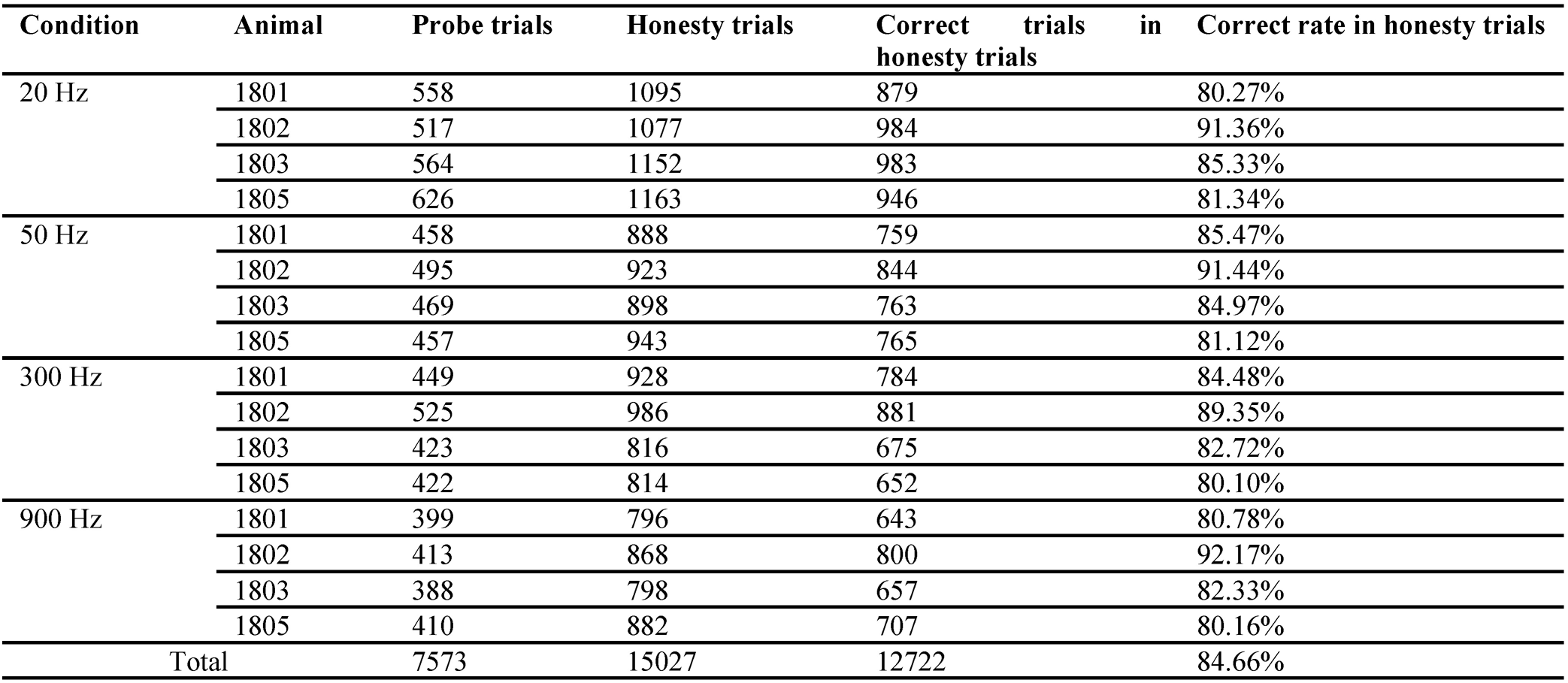
A summary of the data collected in the final “honesty + probe” testing stage.

An analysis of the probe trials obtained during these testing sessions using Probit regression revealed a profound and consistent onset dominance (“precedence effect”) across all animals and all click rates. The results were also remarkably consistent across all animals tested, as can be appreciated in **Figure 2**, which shows the TWFs, computed as Probit weights of each for the eight clicks in our click trains (see Methods), for each of the four animals and at each of the four pulse rates tested. The figure allows us to appreciate the high consistency of the behavior results across all animals in our cohort. The first and 2nd click weights were significantly above zero in all 4 animals for click rates of 50 Hz or less, but for higher click rates of 300 and 900 Hz, significantly non-zero weights beyond the first click are only observed sporadically.

**Figure 2:**
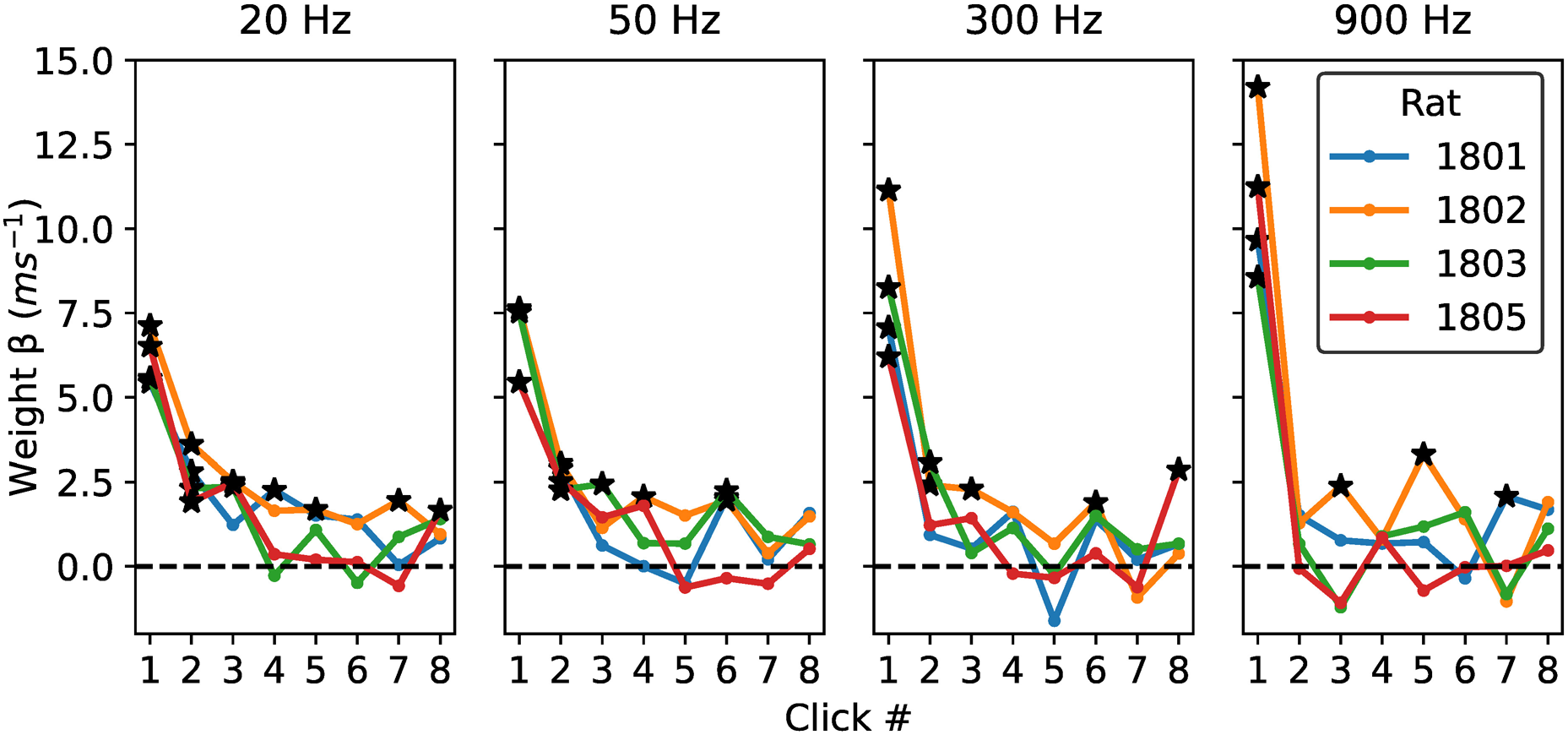
ITD Temporal weighting functions for each individual rat, at the click rates shown above each panel. The x-axes show the individual clicks in the 8-click train. The y-axes show the “temporal weight” of the corresponding click as calculated via Probit regression. Weights which were individually statistically greater than zero based on the *p* values obtained in the Probit regression (*p* < 0.01) are marked with asterisks. Strong onset dominance was seen at all click rates for all rats. The weights of the clicks following the first click increased as the click rate decreased.

It is also noticeable that faster click-rates produced the strongest weighting for the first click, suggesting greater onset weighting at higher pulse rates. This would be consistent with previous reports of similar trends in human listeners (Stecker et al., 2013). In order to estimate whether such differences in weights are statistically significant, we computed bootstrap confidence intervals for the Probit regression weights using classic N-out-of-N resampling. Behavioral data were pooled across all 4 animals, and the set of N trials for each pulse rate, was randomly resampled with replacement to generate a bootstrap sample of size N, for which Probit coefficients were computed in the same way as for the original data set. This random resampling was repeated 1000 times to generate a bootstrap distribution of temporal weights. The resultant bootstrap distributions for the temporal weights are represented graphically as violin plots in **Figure 3**. The full extent to these distributions can be interpreted as a 99.9% confidence interval for the true temporal weight. Note that these do not overlap for click 1 at 20 Hz vs click 1 at 900 Hz, indicating that the observed trend for initial weights to increase with pulse rate is statistically robust. Also note that the bootstrap distributions are entirely above zero at 20 Hz for clicks 1, 2, 3 and 8, while at 900 Hz only the first click had a weight that can be considered significantly greater than zero. This suggests that higher click rates not only produce larger initial weights, but also a more rapid decay of subsequent weights. How complete is this decay? To answer this question we can examine the bootstrap distributions for the later clicks, for example from click 4 onward. Note that, although almost all (15 out of 16) of the bootstrap distributions for clicks 4 to 8 straddle zero, nevertheless the great majority of them (14 out of 16), have medians greater than zero. Such large numbers of above zero medians would not be expected by chance if the later clicks contributed absolutely nothing to the rats’ lateralization percept.

**Figure 3:**
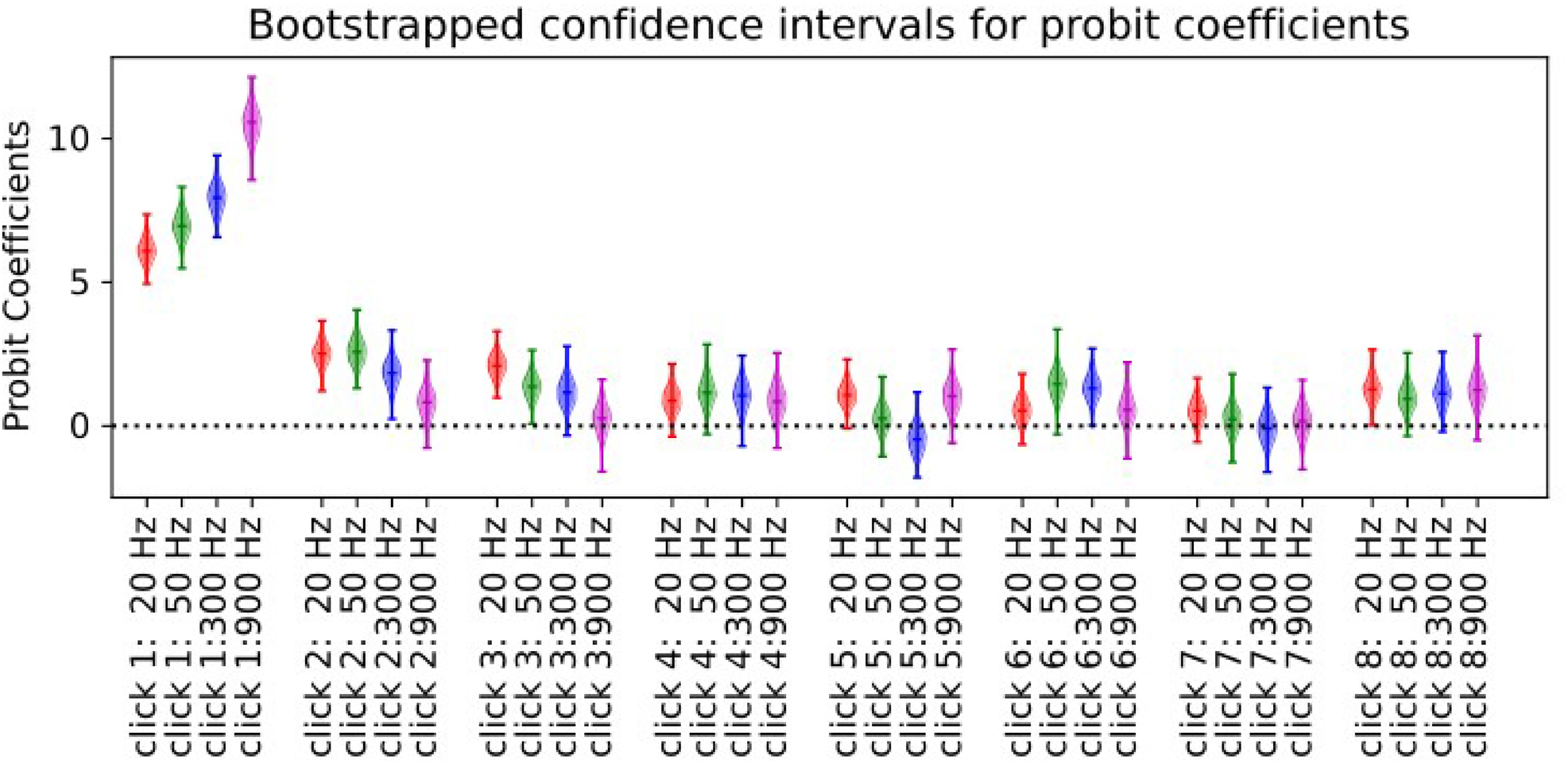
Distribution of Temporal Weights (Probit Coefficients) obtained by resampling the behavioral data. The colored lines show the range and the median values of the coefficient distributions obtained over 1000 bootstrap trials. The spindle shapes display kernel densities of the corresponding distribution: their widths reflect the proportion of bootstrap samples over a given narrow y-range of coefficients in much the same way as the height of a histogram would.

In summary, the behavioral results shown in **Figures 2** and **3** demonstrate that rats temporally weight ITD cues in a manner that is very similar to the temporal weightings previously described for human listeners by Stecker and colleagues (Stecker, Hafter, 2002; Stecker et al., 2013; Stecker, 2018). All four rats in our cohort showed a very strong and highly consistent onset weighting at all tested click rates, and the strength of onset weighting increased significantly at higher click rates.

### 3.2 Channel-wise regression shows only sporadic precedence effect in ECoG signals from the auditory cortex

In total, we recorded electrophysiological data at 12 ECoG electrode placements: 4 placements from the right AC of each of the 4 trained animals in our cohort, and another 3 from the left AC of 3 of the 4 trained animals, plus another 5 recordings from the right AC of 5 untrained animals. At each electrode placement we recorded responses to our sparse TWF stimulus set at two click rates: 300 Hz and 900 Hz, yielding a total of 24 electrophysiology data sets. As described in the methods, in our univariate regression analysis, we aimed to quantify how well changes in LFP response amplitudes could be accounted for by a linear regression model in which each of the four ITDs of our “sparse” ITD stimulus set serve as regressors. This approach does of course rely on the assumption that LFP amplitudes recorded with our ECoG arrays do indeed depend on stimulus ITD, at least to some extent and for some electrode channels. It is of course well documented that single units recorded in auditory cortex with penetrating, high impedance electrodes can be ITD tuned, but one cannot necessarily take it for granted that ITD sensitivity is still observable in ECoG electrode signals recorded from the cortical surface. In **Figure 4** we therefore show example LFPs recorded from a single ECoG channel with two different stimuli, one where all four ITDs were -0.164 ms, the other where all four ITDs were +0.164 ms. It is readily apparent that the LFP amplitude is somewhat smaller for the -0.164 ms ITD case. For this particular channel, that difference in LFP amplitude is highly statistically significant (*p*=0.0026, rank sum test) but that channel was selected as an illustrative example to motivate the overall approach, showing a clear and large difference in line with the type of cortical ITD tuning one might expect to see. If we assume that the precedence effect is established early in the pathway, perhaps by cochlear mechanisms or inhibition in the brainstem and midbrain (we will revisit these notions in the discussion), then we would expect the type of ITD tuning illustrated in **Figure 4** to be dominated by the first click in the train. Furthermore, this dominance ought to come through in the multiple regression model designed described in section 2.3.4.1, which quantifies how strongly RMS LFP amplitude depends on the ITD values of each of the four clicks in the stimulus.

**Figure 4:**
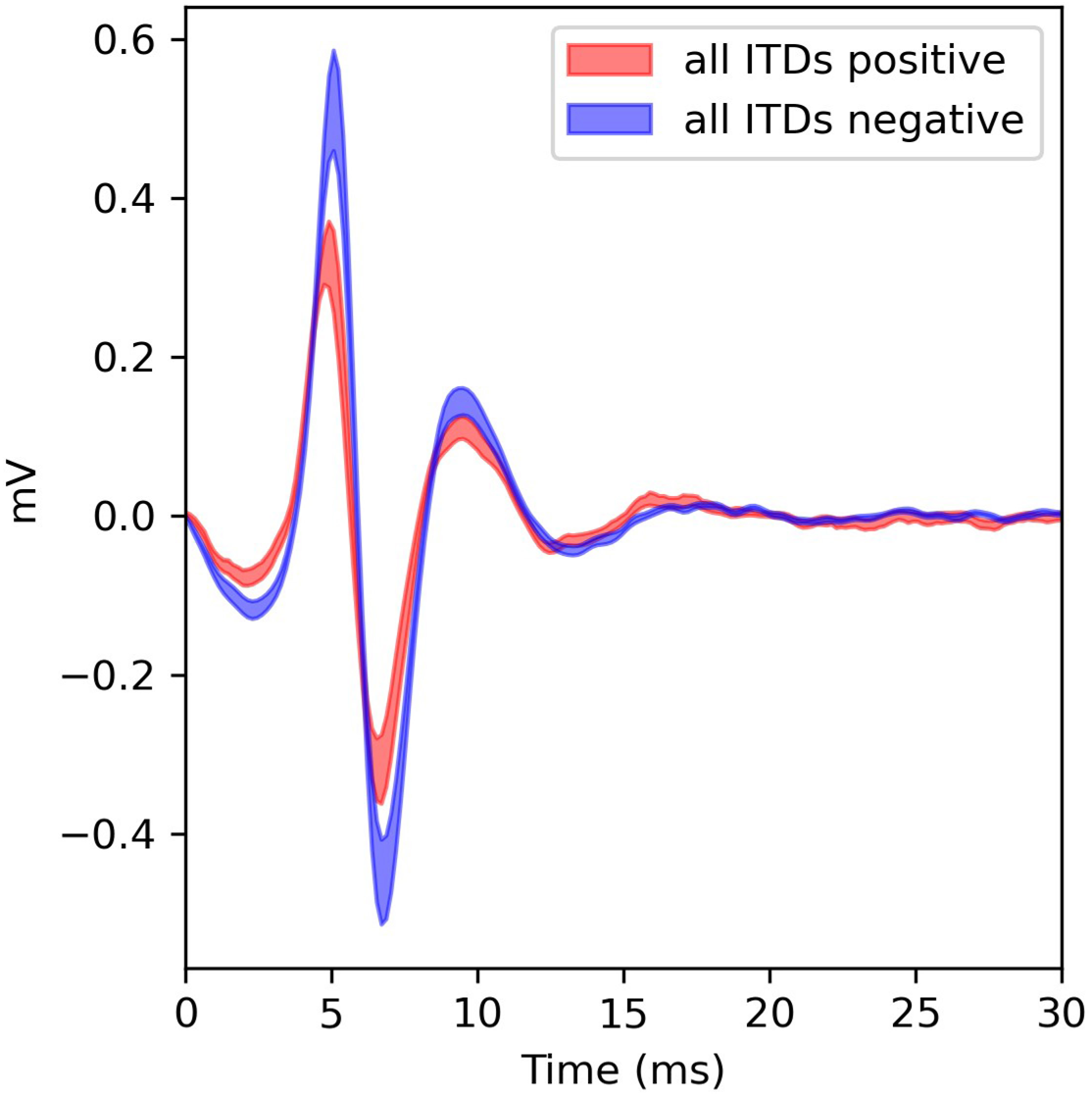
Examples of LFPs recorded at channel 29 during recordings from the left cortex of rat 1803 using the sparse ITD stimulus click train at 300Hz. The lines show mean +/- standard error of the LFP responses recorded when either all four ITDs were positive (red line) or when all were negative (blue line).

However, in contrast to the behavioral experiments, which showed highly consistent and strong weighting of the first pulse, the univariate ECoG results were highly variable and not very consistent in which of the four clicks most strongly shaped the response. Overall, we observed at best only a weak trend for slightly larger weights for the ITD of the first click compared to later clicks in the train, and there was a great deal of heterogeneity in physiological TWFs from animal to animal and from recording site to recording site. Physiological TWF shapes could also vary quite strongly depending on whether the clicks were presented at 300 or 900 Hz. Only for a small subset of ECoG electrode placements did we observe a clear and statistically significant weighting of the first pulse in a majority of channels. An example is shown in **Figure 5A**. For many other electrode placements we saw a much more mixed picture, without a convincing or consistent tendency for the first click to carry the highest weight, as would be expected if neural firing rates consistently reflected the behavioral weighting. The examples in **Figure 5B, 5C and 5D**, illustrate the diversity of physiological TWFs obtained, the strong trend for TWFs to correlate among neighboring channels, and the fact that in these univariate physiological TWFs the largest absolute weights can also often occur on the 2^nd^, 3^rd^ or 4^th^, rather than the 1^st^ click.

**Figure 5:**
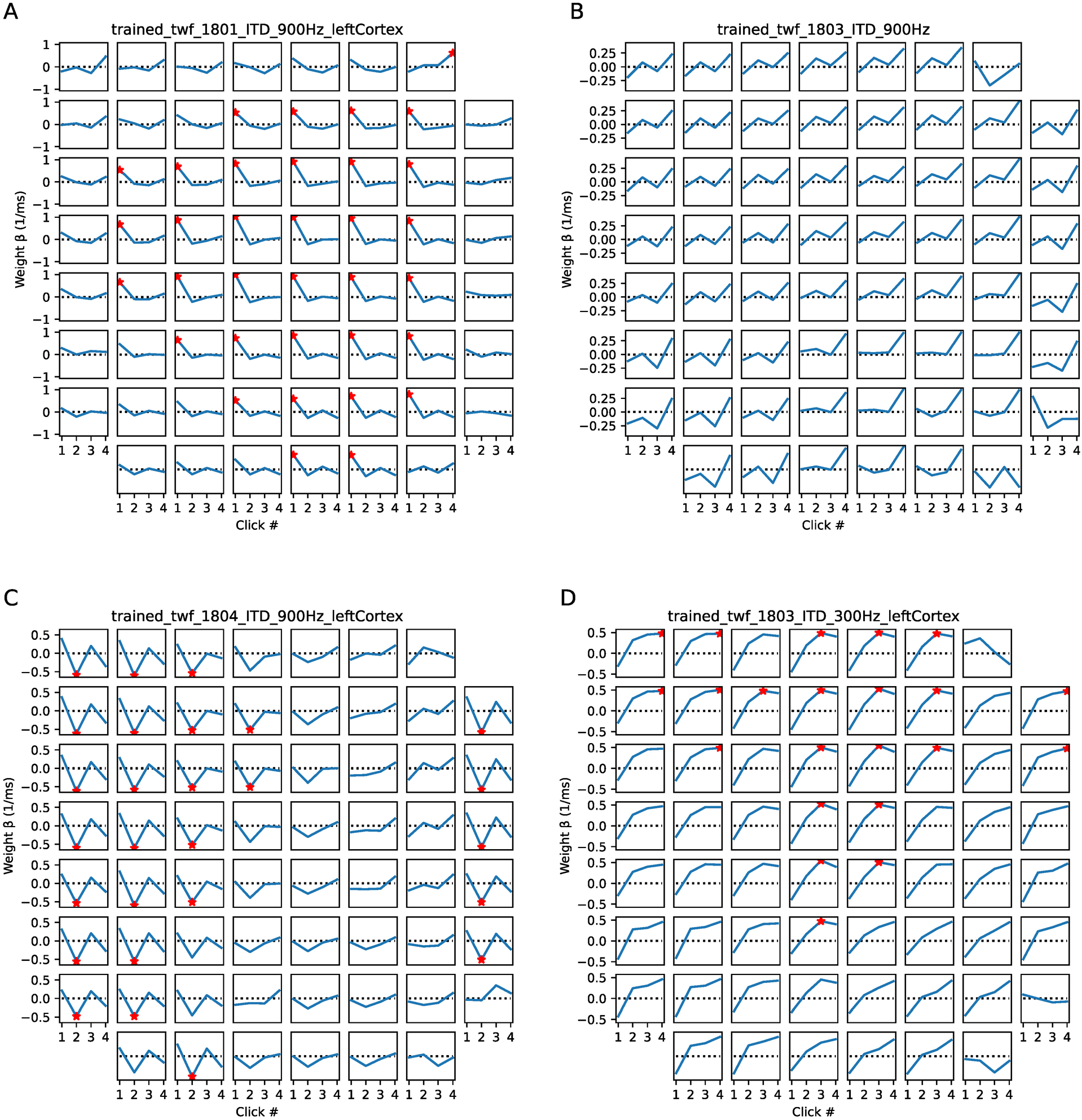
Examples of physiological TWFs obtained by univariate, linear regression analysis of responses recorded with the ECoG array placed on the auditory cortex. In each subplot, the x-axis shows the individual click in the 4-click train and the y-axis shows the beta value (“temporal weight”), in units of proportion change in normalized response per ms ITD, computed with Ordinary Least Squares regression. Each subplot represents the physiological TWF obtained by analyzing the responses recorded at a different ECoG array position and pulse rate. The examples are chosen to be representative of the very diverse results obtained. Red asterisks (*) mark regression weights that were individually significantly different from zero at *p* < 0.05 (not corrected for multiple comparisons). Some physiological TWFs showed strong and significant onset weighting (e.g. most channels in A) but many others did not, and it was not uncommon for temporal weights other than the first to be the largest absolute significant value.

To provide an overview of our complete dataset of 12 electrode placements across the 9 animals, we show boxplots in **Figure 6** which give the distributions of temporal weights (absolute regression coefficients |β|). We chose to plot absolute beta values because the sign of the beta depends on whether the recorded neural population at each electrode site happens to have a preference for ipsilateral or contralateral leading ITDs, and the sign is therefore not relevant to the question of whether the first or second click in the stimulus has a stronger influence on the amplitude of the response. Boxplots are shown separately for the 300 and 900 Hz click rate conditions, and we show both the full distribution of all regression weights and the distributions containing only weights which were significantly different from zero at *p* < 0.05. Our data set comprises a total of 4 coefficients (one for each click in the 4 click train) for each of the 61 electrode channels for each of the 12 electrode placements, yielding a total of a total of 2928 coefficients at each of the two click rates tested. Of these, 147 (5%) were individually significant at alpha 0.05 at 300 Hz, and 228 (7%) were significant at 900 Hz. (Note that one would expect *a priori* that the proportion of significant regression weights obtained in this analysis is likely to be small).

**Figure 6:**
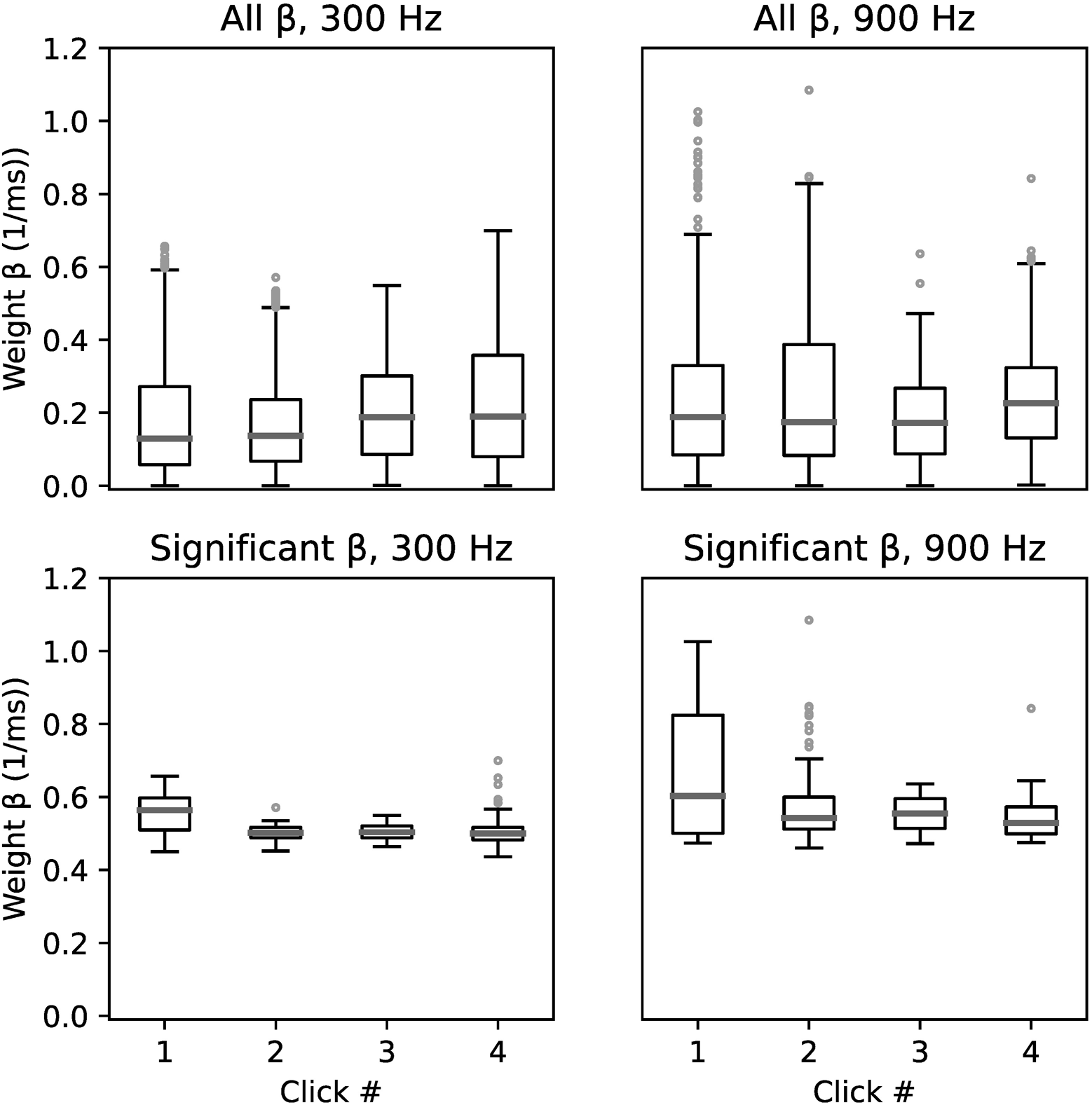
Boxplots showing the distributions of absolute weights which were significantly different from zero (*p* < 0.05) pooled across all ECoG recordings from all four rats at 300 Hz and 900 Hz click rates. The x-axis indicates the individual click in the 4-click train and the y-axis shows the absolute beta value (“temporal weight”) obtained from the Ordinary Least Squares regression. There is a trend for the median absolute weightings on the first click to be larger than for the other clicks, but the trend appears to be very modest when compared to the behavioral TWFs seen in Figure 2, where the weights on the first click were an order of magnitude larger than those seen on the later clicks. The statistical significance of that trend is doubtful and very difficult to assess accurately given the non-normal nature and the nested statistical dependencies of the individual observations.

The median absolute beta values for the first, second, third, and fourth clicks were similar in all conditions. For the distributions that excluded non-significant betas, there was a weak trend for median absolute weight at onset to be larger than the other weights, but this trend is surprisingly weak compared to the robust behavioral onset weighting shown in **Figures 2** and **3**). Note in **Figure 5** that TWFs seen at neighboring recording sites are often highly correlated. Therefore, regression weights obtained from neighboring sites cannot be treated as statistically independent observations, which makes it very difficult to judge whether the observed trends in the ECoG derived weight distributions are statistically significant. In any event, the “effect size” of the weak trend seen in **Figure 6** appears too small, and the neural response patterns are too variable, to provide a convincing physiological correlate of the very strong and consistent onset weighting seen in the behavioral results.

### 3.3 Multivariate decoding shows strong precedence effect in ECoG signals from the auditory cortex

We have just seen that the univariate (channel-by-channel) analysis revealed only a weak trend toward onset weighting in the physiological TWFs obtained by regression analysis, and many individual recording sites showed the strongest weighting for clicks other than the first. In contrast, a population decoding approach revealed very strong and highly significant onset weightings in the physiological responses which closely mirrored the behavioral results. When analyzing the average RMS activity observed within the first 30 ms after click train onset, and pooling RMS values over multiple channels (**Figure 7A**), the ITD decoding estimate of the first click was markedly higher than that for the other clicks. The decoding of the first three clicks yielded decoding values that were significantly different from zero (first click: *p* < 0.001, Z = 3.945; second click: *p* = 0.019, Z = 2.346; third click: *p* = 0.042, Z = 2.033), while the decoding of the fourth click was not significant (*p* = 0.5663). The decoding of the first click was significantly higher than the decoding of all remaining clicks (all *p* < 0.003), and the decoding of the second click was significantly higher than the decoding of the fourth click (*p* = 0.019), while there were no significant differences in decoding between the remaining clicks (all *p* > 0.05). The decoding of the first click was also significant when both click rates were analyzed separately (300 Hz: *p* = 0.009, Z = 2.599; 900 Hz: *p* = 0.003, Z = 2.934; **Figure 7C and D**). Overall, based on neural responses to the click trains pooled from multiple ECoG channels, the ITD of the first click (and, to a lesser extent, of the subsequent two clicks) could be decoded reliably from the cortical population response, and these multivariate decoding results dovetail with the behavioral results which show that the rats base their sound direction judgments predominantly on the first click.

**Figure 7:**
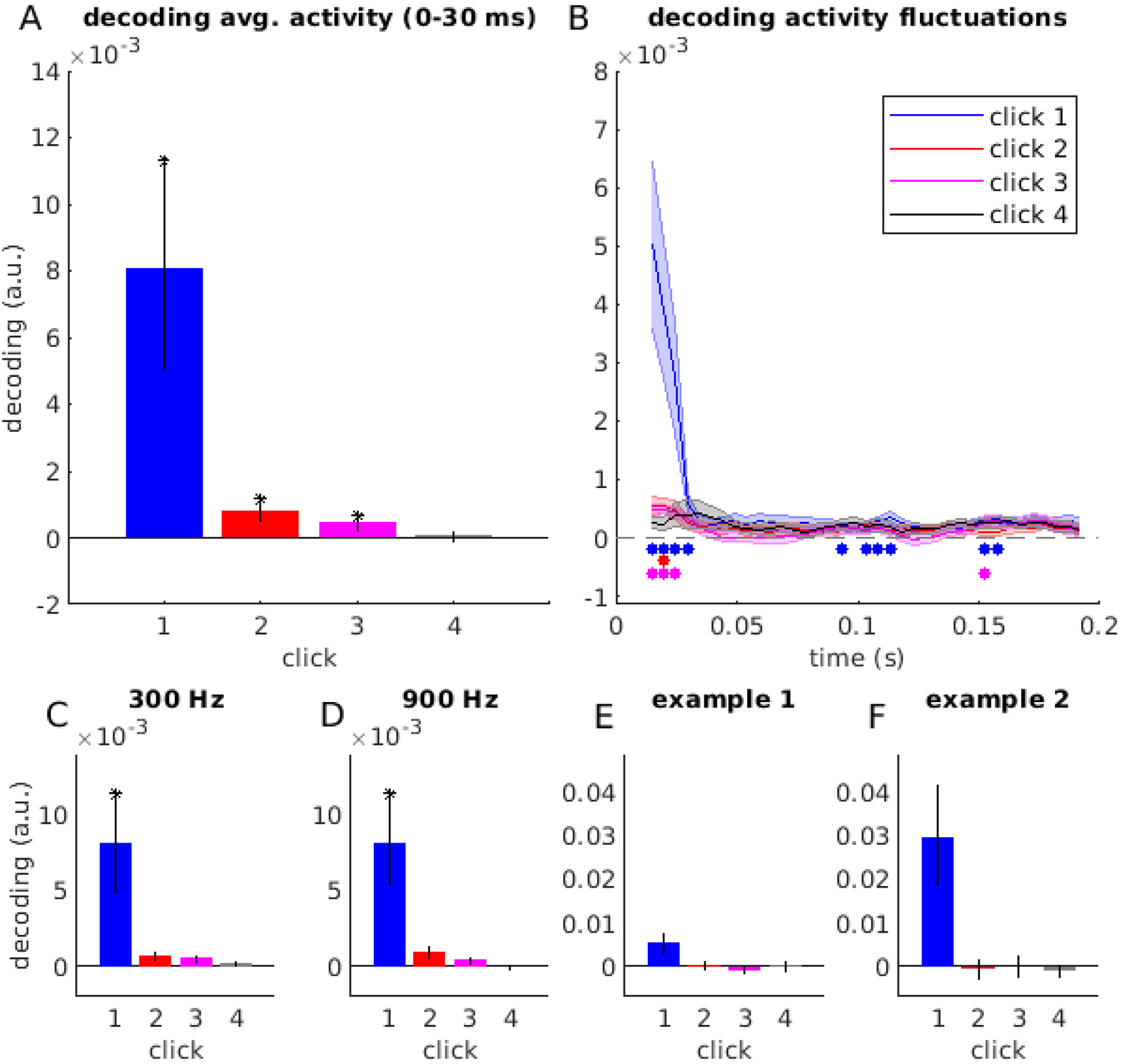
(A) ITD decoding based on average RMS activity between 0-30 ms relative to click burst onset. Error bars denote SEMs across ECoG array placements. Asterisk indicates statistical significance (*p* < 0.05, Bonferroni-corrected). (B) ITD decoding time-series based on activity fluctuations in each 30-ms-long sliding time window. Shaded areas denote SEMs across placements. Filled circles indicate statistical significance (*p* < 0.01, FDR-corrected). (C and D) ITD decoding based on average RMS activity between 0-30 ms relative to click burst onset, separately for 300 Hz and 900 Hz click rates. Error bars denote SEMs across ECoG array placements. Asterisk indicates statistical significance (*p* < 0.05, uncorrected). (E and F) Examples of ITD decoding based on average RMS activity between 0-30 ms relative to click burst onset, separately for two individual penetrations. Example 1 corresponds to Figure 5A; example 2 corresponds to Figure 5B. Error bars denote 99% confidence intervals.

In a further analysis, we investigated the decoding dynamics in a longer time window, ranging from 0 to 200 ms relative to click train onset (**Figure 7B**). In this analysis, rather than using time averaged activity to decode ITD, we used a sliding time window approach, in which we decoded ITD based on RMS fluctuations within 30 ms long time windows (with a time step of 5 ms). Decoding based on activity fluctuations (rather than average activity) makes this analysis especially sensitive to short-lived transients in neural activity (Wolff et al., 2020). This analysis revealed that the ITD of the first click pair could be decoded based on early neural responses, corresponding to time windows centered at 15-30 ms following click train onset (*p*_FDR_ < 0.01, corrected across time windows), but also based on later neural responses (95-115 ms and 155-160 ms following click train onset; *p*_FDR_ < 0.01). Similarly to the decoding analysis based on average activity, in this analysis we also observed weaker but significant decoding of the ITDs of the second and third clicks (click pair 2: 20 ms; click pair 3: 15-25 ms and 155 ms following click train onset; all *p*_FDR_ < 0.01, corrected across time windows; click pair 4: no significant decoding time windows), suggesting that brief neural transients to single click pairs might be sensitive to individual ITDs at different latencies. However, in this analysis too the peak ITD decoding value for the first click was significantly (all *p* < 0.001) larger than those for the later clicks, by approximately an order of magnitude. The decoding results in **Figure 7B** thus closely mirror the behavioral results shown in **Figure 2**.

In **Figure 7E and F**, we plot the individual decoding estimates for the same two examples as plotted in **Figure 5A and B**, respectively. While for both examples the decoding of the first click ITD is better than of the remaining clicks, this trend is even more pronounced for the TWFs where individual channels were heterogeneous and showed no obvious preference for the first click in the univariate analysis (example 2; **Figure 5B**) than for the electrode placement in which univariate TWFs often showed a similar onset preference across the majority of channels (example 1; **Figure 5A**). This is likely due to the fact that we used a Mahalanobis distance metric to estimate ITD decoding, whereby the influence of each channel is scaled by the overall covariance across channels. Therefore, in the case of example 1, where most channels show a similar TWF pattern, the decoding relies on fewer response features than in the case of example 2, where different channels can have a unique contribution to decoding.

## 4 Discussion

### 4.1 Principal findings

To the best of our knowledge, this is the first study to measure binaural TWFs behaviorally and physiologically in a species other than humans. We were able to show that laboratory rats strongly onset weight binaural cues, basing lateralization judgments almost entirely on the first click in a click train. Their perception of such stimuli thus closely resembles that reported for human listeners. Note also that prior human psychoacoustic studies reported somewhat smaller weights on the first click when click rates were fairly low (20 or 100 Hz, inter-click intervals (ICI) of 5-10 ms) compared to those obtained with faster rates (500 Hz; ICIs of 2 ms) (Stecker et al., 2013). Our rat data exhibit a similar trend, with the weight of the first click in a train increasing at higher click rates.

Having thus validated the rat as a good model for human binaural hearing, we were able to follow this up with a physiological examination of the encoding of binaural cue values in cortical population responses recorded with ECoG arrays. Perhaps surprisingly, neural responses obtained at individual recording sites often failed to show a robust onset dominance in the temporal weighting of ITDs, but a multivariate analysis of single trial population responses revealed a neural encoding of ITDs that mirrored the animals’ behavioral responses. This observation has important implications for future studies which may wish to use physiological tests on rats to probe factors governing binaural hearing performance, for example in order to develop improved neuroprosthetic devices. Studies focusing heavily on reading out cortical responses one neuron or one unit or one recording site at a time could be missing out on important correlates of psychoacoustic performance that are only readily apparent at the level of a population coding analysis.

### 4.2 Neural decoding in the auditory cortex as a correlate of the precedence effect

As discussed in (Brown et al., 2015), although the basic features of the precedence effect have been described for more than half a century, its biological mechanisms remain controversial. Electrophysiological recordings from central auditory structures have demonstrated reduced neural responses to “simulated echoes”, that is, pairs of pulses, similar to the first two clicks in the click trains used in this study (Yin, 1994; Litovsky, Yin, 1998; Tollin et al., 2004). It is natural to assume that a reduction in strength of auditory midbrain responses to the second click in a pair might lead to a corresponding down-weighting of that second click “further downstream”, as the neural representations of the clicks are fused into a unified percept of a single source with a (usually not consciously perceived) echo and with a single perceived source location. Rather than challenging this assumption, much previous work has instead focused on discussing the physiological mechanisms that are likely to be responsible for the reduction of the response to the second click. The two main mechanisms considered were synaptic inhibition (Burger, Pollak, 2001; Pecka et al., 2007; Xia et al., 2010) or more peripheral processes involving cochlear mechanics (Hartung, Trahiotis, 2001; Bianchi et al., 2013). Central inhibition and cochlear mechanics could in principle both contribute to the physiological mechanisms which underpin the precedence effect, but recent evidence suggests that cochlear mechanics are unlikely to be the major determinant, given that focal lesions (Litovsky et al., 2002) and pharmacological manipulations (Burger, Pollak, 2001; Pecka et al., 2007) of auditory midbrain structures which leave the cochlea untouched nevertheless alter the precedence effect, and given that psychoacoustic signatures of a precedence effect have been observed even in cochlear implant patients in whom cochlear mechanics have been completely bypassed by neuroprosthetic stimulation (Brown et al., 2015).

Our approach to studying possible physiological bases for the precedence effect differs from previous work in a number of important ways. For example, previous physiological studies commonly measured responses to click pairs delivered in rapid succession in the inferior colliculus and interpreted an observed suppression of the lagging click as a putative correlate of the precedence effect. However, potential echos can arrive very quickly if reflective surfaces are slow. The highest pulse rate tested with our TWF paradigm is 900 Hz, corresponding to inter-pulse intervals of 1.1 ms, which is much too fast for the IC to produce individual responses locked to individual clicks (Schnupp et al., 2015), so here we asked how strongly the neural response as a whole is influenced by the ITD of a given click (univariate, channel-wise regression analysis), and to what extent the response might be “decoded” to reveal the ITD of each click (multivariate analysis). Also, our study looks at neural responses in cortex rather than IC. In this context it is worth mentioning that a previous non-invasive study on human volunteers had described parallels between location judgments under conditions that required echo suppression and auditory evoked potentials (Liebenthal, Pratt, 1999).

Our investigation of the neural correlates of TWFs using channel-wise regression of individual channels produced very variable and inconsistent results, with some recording sites showing the expected largest weights for the first click in the train, but many others showing highly unexpected TWF patterns, with the largest absolute weight placed on the 2^nd^ or 3^rd^ click, in the middle of the 4-click train (cf **Figures 5C and D**). The frequent, large and significant weightings of later clicks is highly surprising because it implies that, at the level of individual groups of cortical neurons, neural TWFs can differ substantially from the behavioral TWF, and place the largest weight away from the stimulus onset. This phenomenon was so common that, overall, there was only a small trend for the first click in the series to have on average (median) the largest absolute weight. This surprising finding led us to hypothesize that, at the level auditory cortex, a consistent precedence effect may only emerge at the level of neural population responses. To test this hypothesis, we analyzed the ECoG data using recently developed neural population decoding methods. That approach produced results in perfect agreement with the precedence effect demonstrated in the behavioral tests.

The two analysis approaches are conceptually and mathematically distinct, and the observed discrepancies between the two sets of results might be due to several reasons. One plausible explanation is that decoding the activity of a wider population of neurons is needed to observe the precedence effect. The regression analysis was applied here to analyze neural activity on a channel-by-channel basis, with a more localized, smaller set of neurons contributing to the responses within one channel. Conversely, the multivariate decoding method selected channels with a high signal-to-noise ratio, and pooled signals from these channels, integrating activity patterns over a much larger population of neurons. In the multivariate analysis, we also scaled the responses of each channel by their noise covariance, accounting for dependencies between channels. The success of the multivariate pattern analysis that we used for neural decoding suggests the following insights into the neural correlates of the precedence effect: First, pooling neural activity over space (channels) – which can enhance neural decoding accuracy (Grootswagers et al., 2017; Nemrodov et al., 2018) – appears to be necessary to uncover a compelling neural correlate of the precedence effect. Second, pooling neural activity over time (i.e., including temporal transients rather than time-average responses as decoding features) – which highlights dynamic, short-lived neural activity patterns (Wolff et al., 2020) – uncovered weaker but significant decoding of the later clicks in the train, so although the precedence effect strongly dominates the neural population code revealed by this analysis, read-outs of this code at fine temporal resolution nevertheless give access to binaural cue values beyond the very onset of the stimulus. This temporal integration of the signals to combine information from separate populations is also reminiscent of observations by Scholl et al., (2010) that the onset and offset of sound stimuli may be represented by non-overlapping populations of primary auditory cortex neurons.

### 4.3 Neural correlates of the precedence effect in mammalian auditory cortex

Our neural decoding results showed a strong precedence effect in the AC of rats for click train stimuli at both 300 and 900 Hz which nicely paralleled the behavior results, but this correspondence is of course not sufficient to prove that the precedence effect results from cortical population coding. Indeed, despite decades of study, the exact role of AC in spatial hearing remains somewhat unclear, and there may be substantial species differences. Unilateral lesions of AC result in poor performance in localizing brief sound in the contralateral sound field in a variety of species (Cranford et al., 1971; Jenkins, Merzenich, 1984; Kavanagh, Kelly, 1987; Lomber, Malhotra, 2008), while other studies indicate an important role for A1 in recalibrating binaural hearing after periods of partial monaural deprivation (Bajo et al., 2019). However, unilateral lesions on the contralateral AC of rats reportedly did not disrupt the sound localization performance of rats (Kelly, 1980). Also, how difficulties in localizing sources within one hemisphere relate to the sound lateralization ability across the midline has not been investigated in detail. There are also previous studies documenting ITD sensitivity at the level of AC of rats (Kelly, Phillips, 1991; Tsytsarev et al., 2009), as well as studies on humans and cats which suggest that an intact AC may be necessary for the precedence effect (Cranford, Oberholtzer, 1976; Cornelisse, Kelly, 1987). The fact that our ECoG data reveal fairly widespread, significant ITD sensitivity cues in LFP responses is therefore unsurprising. However, our finding that population decoding is necessary to reveal a strong precedence effect which mirrors behavioral observations is very novel and points to the unexpected key role that cortex may play in transforming physical binaural cue values into integrated percepts.

## Acknowledgements

Not applicable.

## Authors’ contributions

KL designed and conducted experiments, analyzed and interpreted data, drafted and revised manuscript. RA analyzed and interpreted data, drafted and revised manuscript. CC and AM conducted experiments. JS designed experiments, analyzed and interpreted data, revised manuscript. All authors read and approved the final manuscript.

## Declarations

### Ethics approval and consent to participate

All experimental protocols were assessed and approved by the Experimental Animals Research Ethics Subcommittee at the City University of Hong Kong, and performed under license by the Department of Health of Hong Kong [Ref. No.: (18-9) in DH/SHS/8/2/5 Pt.3].

### Consent for publication

Not applicable.

